# QTL Analysis of Domestication Syndrome in Zombi Pea (*Vigna vexillata*), an Underutilized Legume Crop

**DOI:** 10.1101/353029

**Authors:** Sujinna Dachapak, Norihiko Tomooka, Prakit Somta, Ken Naito, Akito Kaga, Peerasak Srinives

## Abstract

Zombi pea (*Vigna vexillata* (L.) A. Rich) is an underutilized crop belonging to the genus *Vigna*. Two domesticated forms of zombi pea are cultivated as crop plants; seed and tuber forms. The cultivated seed form is present in Africa, while the cultivated tuber form is present in a very limited part of Asia. Genetics of domestication have been investigated in most of cultivated *Vigna* crops by means of quantitative trait locus (QTL) mapping. In this study, we investigated genetics of domestication in zombi pea by QTL analysis using an F_2_ population of 139 plants derived from a cross between cultivated tuber form of *V. vexillata* (JP235863) and wild *V. vexillata* (AusTRCF66514). A linkage map with 11 linkage groups was constructed from this F_2_ population using 145 SSR, 117 RAD-seq and 2 morphological markers. Many highly segregation distorted markers were found on LGs 5, 6, 7, 8, 10 and 11. Most of the distorted markers were clustered together and all the markers on LG8 were highly distorted markers. Comparing this *V. vexillata* linkage map with a previous linkage map of *V. vexillata* and linkage maps of other four *Vigna* species demonstrated several macro translocations in *V. vexillata*. QTL analysis for 22 domestication-related traits was investigated by inclusive composite interval mapping in which 37 QTLs were identified for 18 traits; no QTL was detected for 4 traits. Number of QTLs detected in each trait ranged from 1 to 5 with an average of only 2.3. Tuber traits were controlled by five QTLs with similar effect locating on different linkage groups. Large-effect QTLs (PVE > 20%) were on LG4 (pod length), LG5 (leaf size and seed thickness), and LG7 (for seed-related traits). Comparison of domestication-related QTLs of the zombi pea with those of cowpea (*Vigna unguiculata*), azuki bean (*Vigna angularis*), mungbean (*Vigna radiata*) and rice bean (*Vigna umbellata*) revealed that there was conservation of some QTLs for seed size, pod size and leaf size between zombi pea and cowpea and that QTLs associated with seed size (weight, length, width and thickness) in each species were clustered on same linkage.

## Introduction

The genus *Vigna* is an important taxon of leguminous plants. This genus comprises more than 100 species that distribute in all major continents including Africa, America, Asia, Australia and Europe. Among those species, as high as nine *Vigna* species are domesticated/cultivated. These domesticated *Vigna* species include azuki bean (*Vigna angularis* (Wild.) Ohwi and Ohashi), black gram (*Vigna mungo* (L.) Hepper), créole bean (*Vigna reflexo-pilosa* Hayata), mungbean (*Vigna radiata* (L.) Wilczek), moth bean (*Vigna aconitifolia* (Jacq.) Maréchal), rice bean (*Vigna umbellata* (Thunb.) Ohwi and Ohashi), cowpea (*Vigna unguiculata* (L.) Walps.), Bambara groundnut (*Vigna subterranea* Verdc.) and zombi pea (*Vigna vexillata* (L.) A. Rich) [1,2]. The former six species are of Asian origin and belong to the same subgenus *Ceratotropis*, while the latter three species are of African origin. Cowpea and Bambara groundnut belong to the subgenus *Vigna*, while zombi pea belongs to the subgenus *Plectrotropis*. These crops are generally cultivated by resource-poor farmers as a sole crop or a component in various cropping systems.

Among the domesticated *Vigna* species, zombi pea is the least known crop. Two forms of zombi pea exists; seed type and tuber (storage root) types. The seed type is believed to be domesticated in Sudan (Africa) [3;4], while the tuber type is believed to be domesticated in Bali and Timor, Indonesia (Asia) [5] and India [6,7]. Genetic diversity analysis in a large set of *V. vexillata* germplasm using simple sequence repeat (SSR; also known as microsatellite) markers revealed clear difference between the two types and suggested that these two types were domesticated from different wild gene pools of *V. vexillata* [4]. The seed type zombi pea is classified as variety *macrosperma*. Possibly, this type is domesticated from wild zombi pea of East Africa [4]. It has erect growth habit and large seed size, and is insensitive to day length and no seed dormancy with some degree of pod dehiscence. The variety *macrosperma* is highly cross-compatible with wild *V. vexillata* [8]. The tuber type zombi pea has not yet been classified as a variety of *V. vexillata*. This type is likely to be domesticated from wild zombi pea of Asia [4]. It has viny growth habit and large seed size, and is sensitive to day length and no seed dormancy with some degree of pod shattering. The tuber type is grown principally for fresh edible tuber roots which contain as high as 15% of proteins [5,6]. This cultivated type has been reported to be cross-incompatible with wild *V. vexillata* and the variety *macrosperma* [8]. The phenotypic difference between the two cultivated types of *V. vexillata* is a result of divergent selection during the evolution of the crop. Morphological and physiological differences that occur during the change from wild plants to cultivated crops is known as “domestication syndrome” [9]. Crop domestication is an accelerated evolutionary process that is the result from human intentional and unintentional selection and natural selections [10]. Domestication-related traits in legume crops include plant architecture (determinate growth habit), gigantism in the consumed plant organs (seed size and/or pod size), reduced seed dispersal (indehiscent pod), loss of seed dormancy and no response to photoperiod [11–13].

Knowledge on genetic basis of crop domestication is useful for identification of beneficial gene(s) in the wild relatives of the cultivated crops that can be used in plant breeding. Moreover, such knowledge can also provide insight into crop evolution and agriculture, such that the case of genetic analysis of indehiscent pod in soybean by Funastuki [14] which showed crucial role of the trait in the global expansion of soybean cultivation. Generally, genes controlling crop domestication are identified by quantitative trait loci (QTL) analysis. Among crops of the genus *Vigna*, QTL mappings of domestication syndrome traits have been carried out in azuki bean [13,15], mungbean [16], rice bean [17], cowpea/yardlong bean [18–21]. The genetics of domestication of zombi pea is of particular interest because two cultivated forms of this legume have experienced different domestications from wild zombi pea. Seed type zombi pea was domesticated from wild zombi pea of Africa, while tuber type zombi pea was domesticated from wild zombi pea of Asia [4,22]. Information on the genetics of domestication-related traits would be useful for zombi pea improvement programs, and for comparative genome studies among members of the genus *Vigna*. The objectives of this study were to identify QTLs for domestication-related traits in tuber type zombi pea and to compare them with previously reported QTLs for domestication in other *Vigna* species.

## Materials and Methods

An F_2_ population of 139 plants was used in this study. The population was developed from a cross between accession JP235863 and accession AusTRCF66514 (Fig 1). JP235863 is a cultivated zombi pea collected from Bali Island, Indonesia which it is cultivated principally for edible tuber root, while AusTRCF66514 is a wild zombi pea from India. AusTRCF66514 was crossed with JP235863 as female and male parents, respectively, to produce F_1_ hybrid. Only one F_1_seed was obtained and was self-pollinated to generate F_2_ population. One hundred and eighty seven F_2_ seeds together with parental seeds were sown in pots (1 seed per 1 pot) during June to November 2015 under a greenhouse condition at National Agriculture and Food Research Organization, Tsukuba, Japan. One hundred and fifty eight seeds germinated and developed into plants, but 139 of them grew vigorously and set flowers, and were used for DNA analysis and phenotypic data collection.

**Fig 1.**
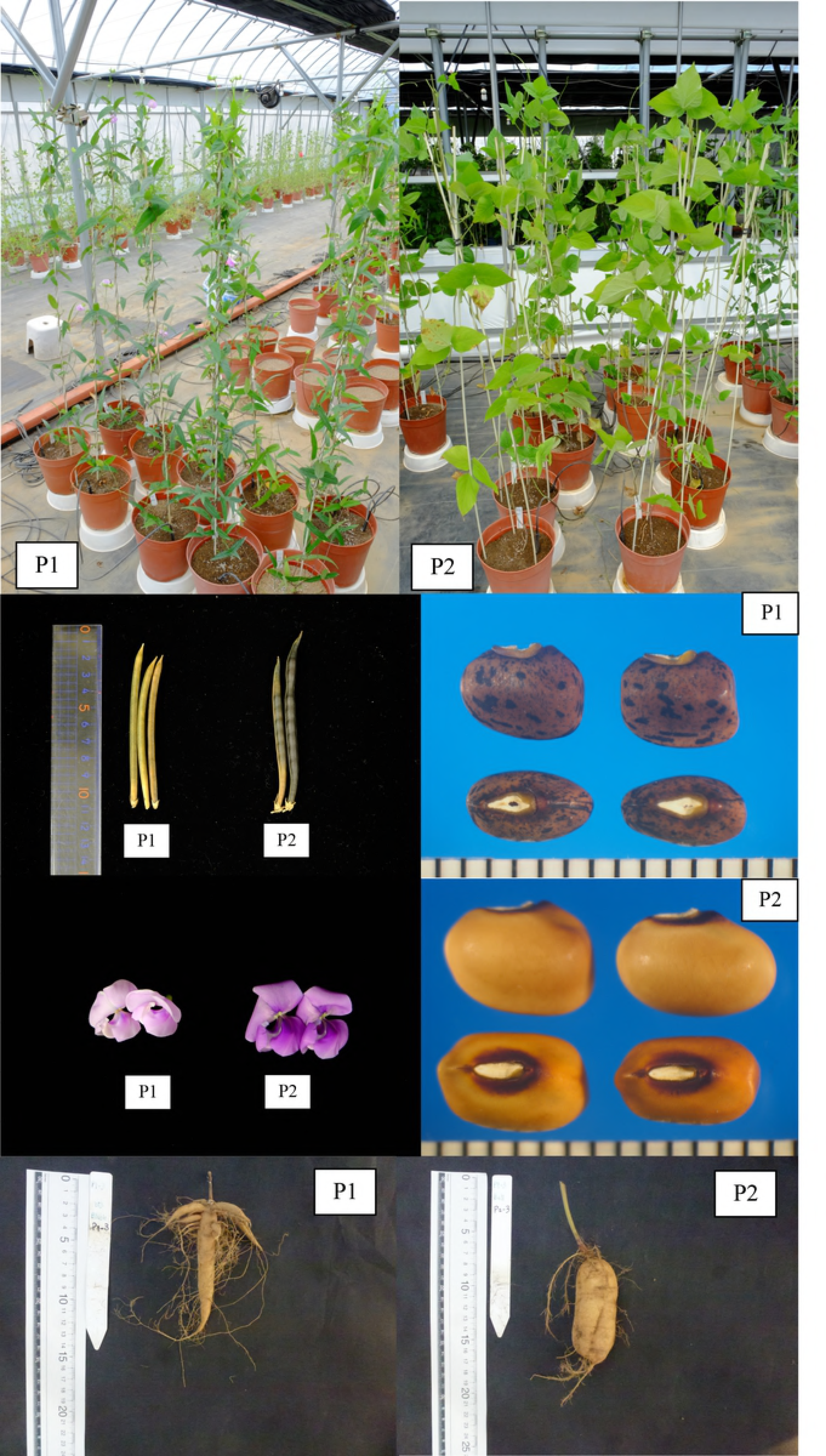
Comparison of morphological characteristics between wild *V. vexillata* accession “AusTRCF66514” and cultivated *V. vexillata* accession “JP235863” used in this study.

### DNA extraction

Total genomic DNA was extracted from young leaves of the parental and the F_2_ plants using a CTAB method described by Lodhi et al. [23]. DNA quality and quantity was checked by comparing with a known concentration ʎ-DNA (50 and 100 ng/μl) on 1% agarose gel. DNA concentration of each plant was adjusted to 5 ng/μl for PCR analysis. The adjusted DNA was also checked for quality using NanoDrop™ 8000 (Thermo Fisher Scientific, Wilmington, U.S.A.).

### Trait measurements

Twenty-two traits related to domestication in *Vigna* crops [13,15–17,20] were measured/evaluated (Table 1). Among those traits, seed color (SDC) and presence of black mottles (SDCBM) on seeds were considered as qualitative trait, while the others were considered as quantitative trait. Seedling traits, primary leaf width (LFPW) and primary leaf length (LFPL) were recorded when the first trifoliate was appeared. Stem thickness (STT), 10^th^ node length (STL10), maximum leaf length (LFML) and maximum leaf width (LFMW) were collected when eleventh trifoliate leaf was fully developed. Seed-related traits including number of seeds per pod (SDNPPD) was an average from ten pods, seed width (SDW), seed length (SDL) and seed thickness (SDT) was the average from ten seeds while 100-seed weight (SD100WT) was recorded from 100 seeds of each plant. The number of days from planting to first flowering (FLD) was recorded. Number of days to first mature pod (PDDM) chosen from first flower that had been developed to mature pod. After harvesting all pods, ten pods from each plant were used to measure/recorded for pod width (PDW), pod length (PDL), pod dehiscence (PDT) or number of twist along the length of pod (NTWP). PDT was recorded after the pods were kept in hot air oven at 40°C for 24 hours. Tuber traits (NTB, TBWT, TBW and TBL) were collected at 260 days after planting.

**Table 1. Domestication-related traits determined in F_2_ population derived from cross between cultivated (Bali) and wild (India) accessions.**

### SSR marker analysis

A total of 1,876 SSR primer pairs (827 pairs from azuki bean [24,25], 562 pairs from cowpea [19,26], 318 pairs from mungbean [27–29], and 169 pairs from common bean [30–32]) were screened for polymorphism between the wild and cultivated parents. PCR amplification and fragment analysis for the primers from azuki bean, cowpea, and common bean were performed following the method described by Marubodee et al. [33], while for the primers from mungbean were the same as described by Somta et al. [34]

### RAD-seq analysis

RAD-seq analysis was conducted following a modified protocol described by Peterson et al. [35]. Genomic DNA of parents and each F_2_ plants was digested by double-digest RAD library method. In brief, 10 ng of genomic DNA was digested with EcoRI and BgIII (New England Biolabs, Ipswichs, MA, USA), and purified. Each adaptor with unique 4-8 bp index was ligated to digested DNA samples. The adaptor sequence was as follows: TruSeq_EcoRI_adaptor 1 = A*A*TTGAGATCGGAAGAGCACACGTCTGAACTCCAGTC*A*C TruSeq_EcoRI_adaptor 2 = G*T*CAAGTTTCACAGCTCTTCCGATC*T*C (* = variable index sequences to identify the individual DNA sample). BglII adaptor =A*A*TGATACGGCGACCACCGAGATCTACACTCTTTCCCTACACGACGCTCTT*C*C TruSeq_BglII_adaptor 2=G*A*TCGGAAGAGCTGTGCAGA*C*T. The adaptors were ligated at 37°C overnight in 10 μl volume which contained 1 μl of 10× NEB buffer 2, 0.1 μl 0f 100 × BSA (New England Biolabs), 0.4 μl of 5 uM EcoRI adapter and BgIII adapter, 0.1 μl of 100 mM ATP and 0.5 μl of T4 DNA ligase (Enzymatic, Beverly, MA, USA). The reaction solution was purified with AMPure^®^ XP (Beckham Coulter, CA, USA). Three microliter of the purified DNA was used in PCR amplification in 10 μl volume, containing 1μl of each 10 μM index and TruSeq universal primer, 0.3 μl of KOD-Plus-Neo enzyme and 1 μl of 10× PCR buffer (TOYOBO, Osaka, Japan), 0.6 μl of 25 mM MgSO_4_ and 1 μl of 10 mM dNTP. Thermal cycling was initiated with 94°C step for 2 mins, followed by 20 cycles of 98°C for 10 s, 65°C for 30 s and 68°C for 30 s. The PCR products were pooled and purified again with AMPure^®^ XP. The purified DNA was then loaded to a 2.0% agarose gel and fragments with size of about 320 bp were retrieved using E-Gel^®^ SizeSelect™ (Life Technologies, Carlsbad, CA, USA), the library was sequenced with 51 bp single-end read in one lane of HiSeq2000 (Illumine, San Diego, CA, USA).

### Extraction of RAD-tag and bi-allelic RAD-marker detection

RAD-tag sequences were extracted and bi-allelic RAD-markers were detected using Stacks ver. 1.30 [36] following the procedures described by Marubodee et al. [33]. With the Stacks pipeline, the low quality sequences were filtered to obtain 51bp RAD-tag sequence reads, and classified the RAD-tags sequences into the parental accessions and F_2_ individuals. Then ran denovo_map.pl program where RAD-tags with less than two sequence mismatches were grouped as a stack, parental stacks with less than three mismatches were estimated as those derived from homologous loci and lists of RAD-tag sequences and their count were constructed for each sample. We called genotypes of F_2_ individuals only when stacks have more than five RAD-tags (minimum depth of 5).

### Linkage map construction using SSR ad RAD-seq markers

A genetic linkage map of the F_2_ population was constructed using software JoinMap4.0 [37]. Polymorphic markers showing more than 10% missing genotypic data were excluded from linkage analysis. Chi-square analysis was used to test segregation of the marker loci (1:2:1 for dominant markers and 3:1 for dominant markers). The markers were grouped using minimum logarithm of the odds (LOD) of 3 and recombination frequency of 0.40. Genetic distance between markers in centimorgan (cM) unit was calculated using Kosambi’s mapping function [38].

### QTL analysis

QTLs controlling domestication-related traits were located onto the linkage map by inclusive composite interval mapping (ICIM) method [39] using the software QTL IciMapping 4.1 [40]. Probability value for entering variables in stepwise regression of phenotype on marker variables (PIN) was 0.001. ICIM was performed at every 1 cM. Significant log of odds (LOD) threshold for QTL analysis of each trait was determined by 3,000 permutation test at *P*=0.05.

## Results

### SSR polymorphism

Out of 1,876 SSR primer pairs screened for polymorphism between the parents, 687 pairs (36.6%) were able to amplify DNA of the parents. Primers from azuki bean showed highest amplification rate (51.4%), while those from cowpea showed lowest amplification rate (19.2%). Among the 687 amplifiable primers, only 201 (29.3%) primers showed polymorphism between the parents. Percentage of polymorphic primers ranged from 20.4 (mungbean primers) to 32.7 (azuki bean primers) (S1 Table).

### RAD-seq markers

Illumina sequencing with HiSeq2000 yielded a total of 29,568,103,697 RAD-tag 51-base reads from 578,901,731 raw reads. The number of RAD-tags of parents (AusTRCF66514 and JP235863) and F_2_ individuals was 3,124,320, 2,700,159 and 573,077,252, respectively. The average number of RAD-tags per F_2_ individual was 4,122,857.93. RAD-tags were aligned and clustered into 6,576 stacks. Average fill gaps rated for this 139 F_2_ individuals was 0.747 (max=0.949, min=0). Among these stacks, 479 RAD markers showed homozygote polymorphic genotype between parents. However, among these 479 RAD markers, 362 RAD markers were discarded because they had percentage of missing data more than 10%.

### Linkage map analysis

Finally, 264 markers (145 SSR, 117 RAD and 2 morphological markers) were used for the linkage map construction after discarding markers showing identical F_2_ genotypes, missing data more than 10%. Among these markers, as high as 128 (48.5%) showed segregation distortion. The 264 markers were clustered into 11 linkage groups (LGs 1-11) (Table 2 and Fig 2). The map spanned 704.8 cM in total, with a mean distance between markers of 2.87 cM. The lengths of linkage groups ranged from 30.9 cM (LG10) to 112.7cM (LG2). A gap between markers more than 15 cM presented on only LG11 (between markers DMBSSR192 and CEDG066).

**Table 2. Number of markers and average distance between markers in each linkage group of zombi pea F_2_ population (AusTRCF66514 x JP235863).**

**Fig 2.**
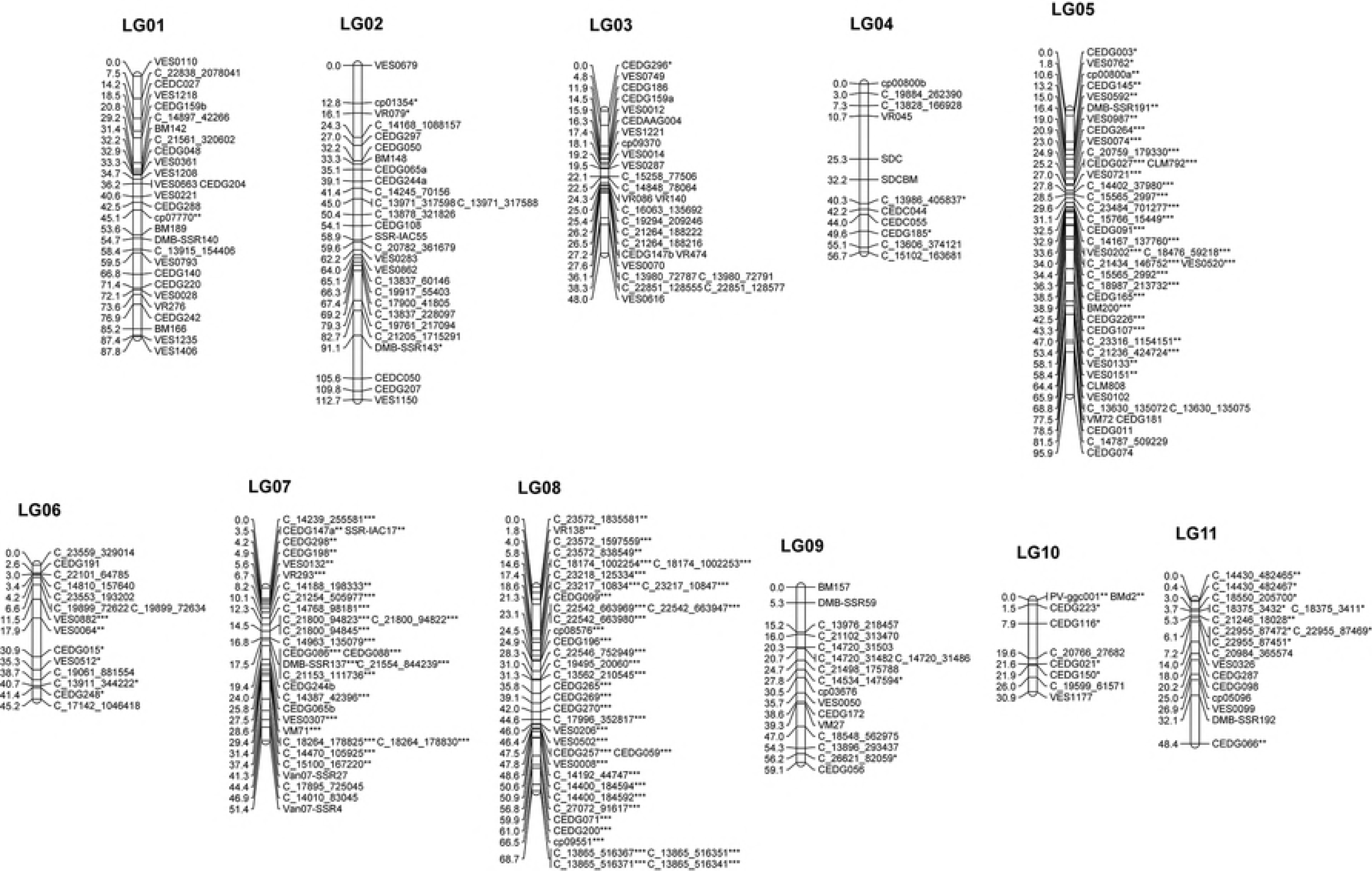
A genetic linkage map of *V. vexillata* developed from F_2_ population of 139 individuals from crossed between AusTRCF66514 and JP235863. Map distance and marker names are shown on the left and right side of the linkage groups, respectively. Markers showing significant deviation from the expected segregation ratio at 0.05, 0.01 and 0.001 probability levels are indicated with *, ** and ***, respectively. Marker names with prefixes C_ are RAD-Seq marker. SSR marker names with prefixes CED, Van07 and VES are from azuki bean. Marker names with prefixes VR and DMB-SSR are from mungbean. Marker names with prefixes cp and VM are from cowpea. Marker names with prefixes BM, BMd, SSR-IAC and PV are from common bean. SDC and SDCBM are morphological markers.

LGs 5, 6, 7, 8, 10 and 11 possessed the several/many distorted markers. The distorted markers clustered together. All the markers on LG8 were distorted markers. LGs 5 and 8 possessed many markers showing severe segregation distortion (*P*< 0.001). All the distorted markers on LG5 showed excessive cultivated parent genotype. Majority of the distorted markers on LGs 6 and 11 and about half of the distorted markers on LG10 showed excessive heterozygous genotype. All the distorted markers on LGs 7 and 8 and about half of the distorted markers on LG10 showed excessive homozygous wild genotype.

Primers CEDG065 and CEDG244 each amplified two loci in which one of the loci of these primers were mapped adjacent to each other and the other loci of these primers also mapped to adjacent to each other. One of the two adjacent pairs of loci was on LG2, while the other adjacent pair of loci was on LG7. This indicated partial genome duplication and translocation of LG2 to LG7 or vice versa.

### Comparative linkage analysis

The linkage map of zombi pea developed in this study were compared with previous linkage maps developed for *Vigna* crops including yardlong bean [19], azuki bean [41], mungbean [16] and rice bean [17] by using common SSR markers (Fig 3). The comparison revealed that in general the linkage groups and orders of the markers among the *Vigna* species were the same or highly similar. Nonetheless, remarkable macro-translocations were found on LG5 and LG7 of our linkage map for *V. vexillata*. About a half of LG5 (middle to bottom) appeared to be a translocation from LG4, while a large portion of LG7 appeared to be translocations from LG4 and LG10. In addition, a small portion of the LG7 appeared to be duplication from LG2, and vice versa. The length of the zombi pea linkage map developed in this study was shorter that the linkage maps of others *Vigna* species.

**Fig 3.**
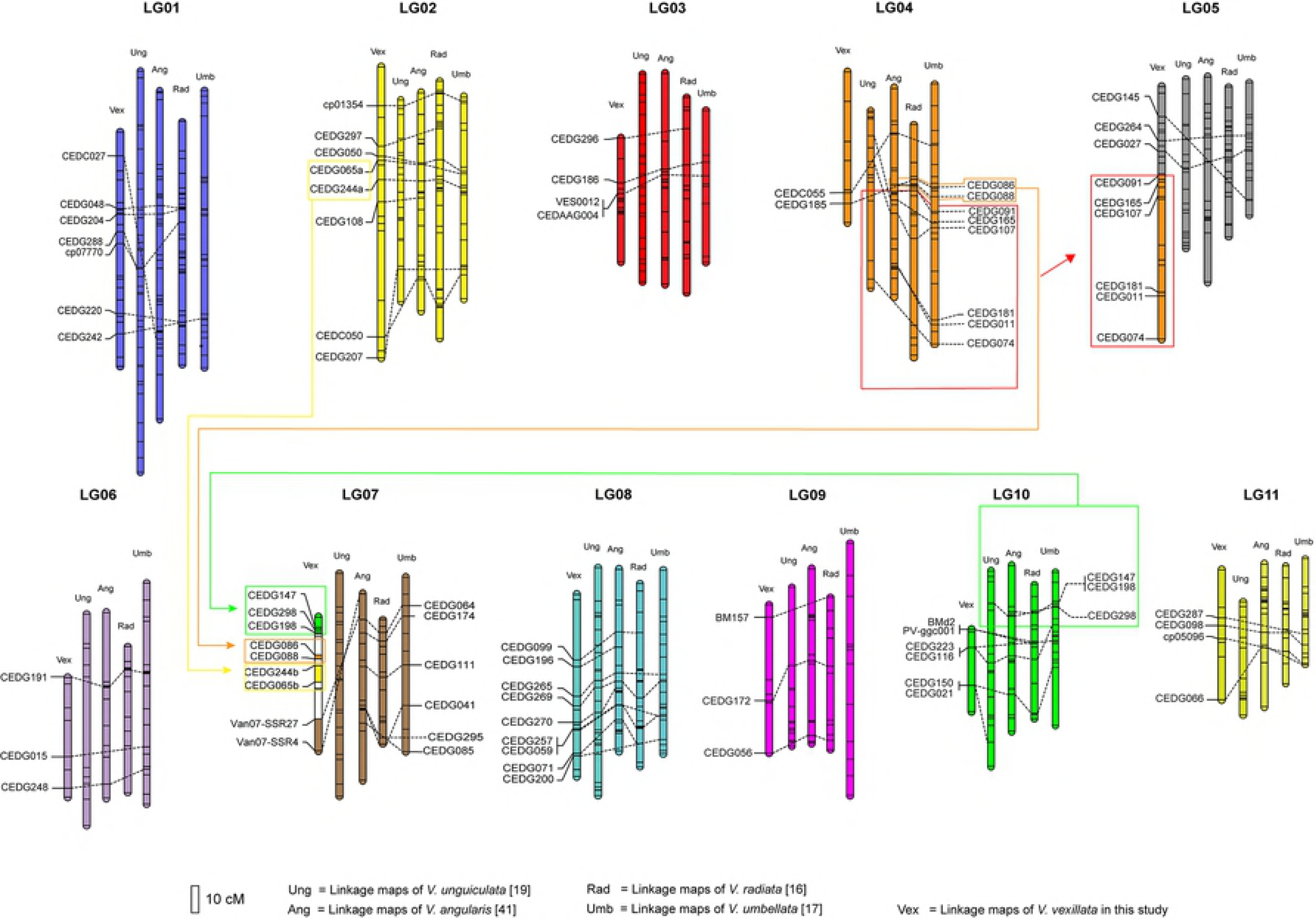
Comparative linkage map among *V. vexillata* (Vex), *V. unguiculata* (Ung), *V. angularis* (Ang), *V. radiata* (Rad) and *V. umbellata* (Umb) based on common SSR markers. Dotted lines connected common SSR markers between linkage maps.

### Variation of domestication-related traits

The parents were clearly different in all the traits recorded. The cultivated zombi pea showed higher value of trait value than the wild zombi pea in all the traits except pod shattering, seeds per pod, tubers per plant, and tuber length. For qualitative traits, the cultivated parent had yellow seed with any mottle, while the wild zombi pea had brown seed with black mottle (Fig 1). The mean, range and standard deviation, and broad-sense heritability of the quantitative traits are shown in Table 3. Mean of the traits of the F_2_ population was between mean of parents for all traits except pod length, tuber length, 10^th^ node length and number of seeds per pod.

**Table 3. Means, standard deviation (S.D.), minimum, maximum and heritability values for the parents and the F_2_ population that derived from a cross between wild (AusTRCF66514) and cultivated (JP235863)**

The measured traits in the F_2_ population showed nearly a normal distribution (Table 3). PDT, SD100WT, SDL, SDW, SDT, PDW, STT, LFPW, LFML, LFMW, TBWT, STL10 and PDDM showed moderate to high heritability (>60%), whereas LFPL, TBW, TBL and FLD showed medium to low heritability (<60%). PDL, NTB and SDNPPD showed heritability lower than 30%.

There was significant and positive correlation (*P*<0.05) between related traits in F_2_ population, such as between seed-related traits (SD100WT, SDL, SDW and SDT), between pod width and seed-related traits (SD100WT, SDL, SDW and SDT), between SDNPPD and PDL, between LFPW and LFMW, between TBW and TBWT (S2 Table). Negative correlation was found between TBW, TBWT and SDNPPD, STL10 and LFML and NTB, TBL and FLD (S2 Table). In general, correlation between related traits was moderate to high (> 0.50), while correlation between non-related traits was low or none.

### QTLs for domestication-related traits

The results of the QTL analysis for each trait in F_2_ population are shown in Table 4. Out of 22 traits, ICIM did not find any QTL for four traits. Among those 18 traits which QTLs were identified, six traits each had only one QTL.

**Table 4. QTLs-related domestication detected in F_2_ from cross between wild *V. vexillata* (AusTRCF66514) and cultivated *V. vexillata* (JP235863).**

**Pod dehiscence** (PDT). Both cultivated and wild parents showed small difference in number of twists along the pod (Table 3). In the F_2_ population, there was little variation of the trait. As a result, no QTL was detected for this trait.

**Increase in organ size** Seed size, pod size, stem thickness, leaf size and tuber-related traits were considered.

**Seed size** (SD100WT, SDL, SDW and SDT): One to three QTLs for traits related to seed size were located on LGs 2, 5, 7 and 8. At all QTLs detected, alleles from the cultivated parent increase the seed size. The QTLs with the largest phenotypic effects for all four traits were located on LG7. (28.19%, 32.60% and 24.02% for seed weight, seed length and seed width, respectively).

**Pod size** (PDL and PDW): The cultivated zombi pea had larger pod size than the wild zombi pea. Three QTLs for each PDL and PDW traits were detected on LGs 1, 2, 4, 5 and 7. The QTLs with the highest contributions to these traits (PVE = 20.65% and 18.11%) were found on LGs 4 and 2. QTLs for both PDL and PDW on LGs 5 and 7 were located close to QTLs for seed size traits (Fig 4).

**Fig 4.**
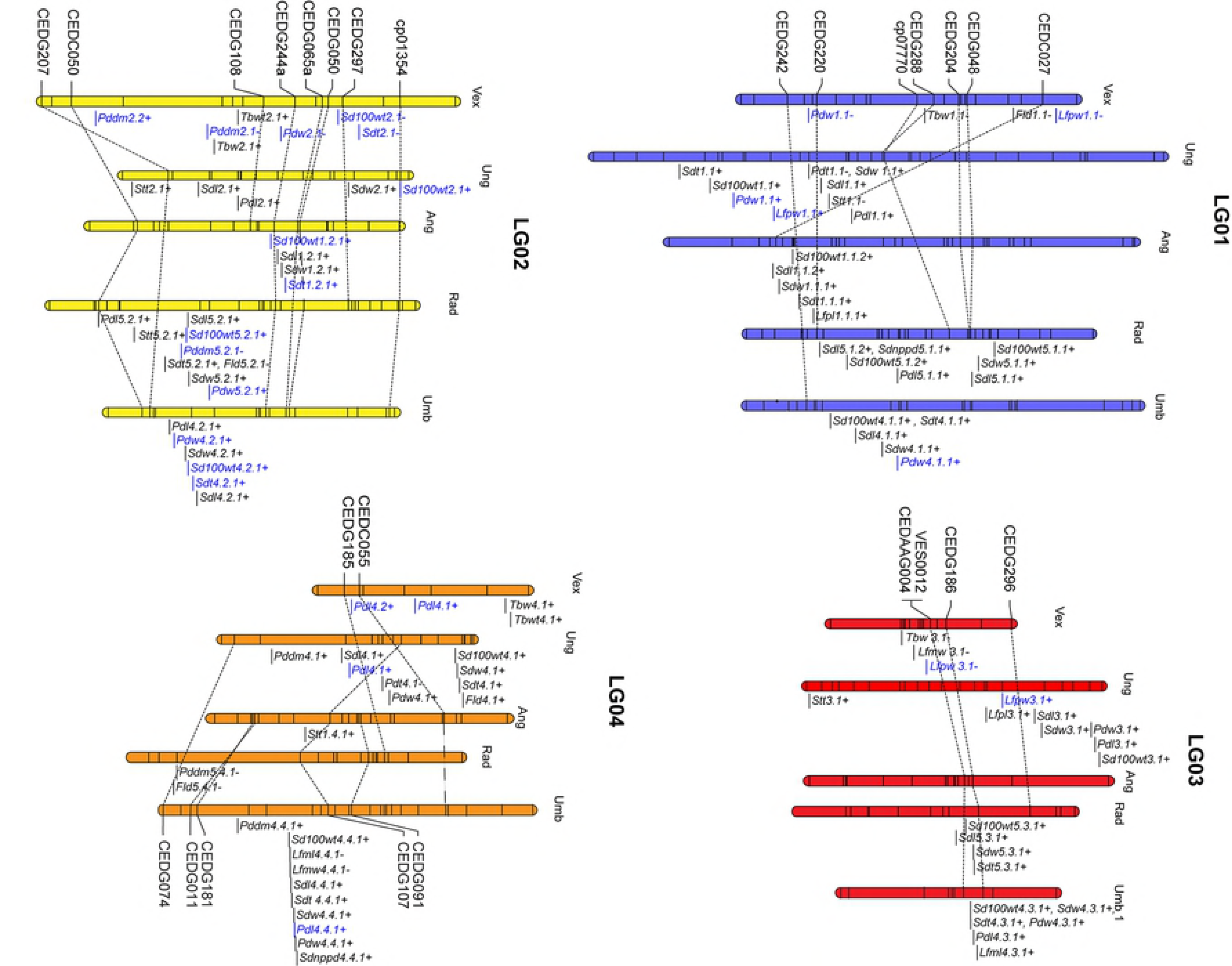
QTLs detected for domestication-related traits in *V. vexillata* F_2_ population of the cross between wild (AusTRCF66514) and cultivated (JP235863) accessions. The effect of the cultivated parent is indicated after each name. Explanation of trait abbreviation is shown in Table 1. QTL names in blue color are QTLs that are also identified on same linkage group(s) of other *Vigna* species. Bold QTLs names are common QTLs between/among species.

**Stem thickness** (STT): Only one minor QTL (PVE = 11.25%) was detected on LG7 for stem thickness which was influenced by alleles from the cultivated parent that has thicker stem than the wild zombi pea.

**Leaf size** (LFPL, LFPW, LFML and LFMW): Only one QTL was detected on LG8 for LFPL. Three QTLs for LFPW were detected on LGs 1, 3 and 5. One QTL was detected on LG5 for LFML. Two QTLs for LFMW were detected on LGs 3 and 5. The QTLs on LG5 for LFPW, LFML and LFMW showed highest PVE and were located very close to each other and likely to be the same locus. Similarly, the QTLs located on LG3 for LFPW and LFMW were located not far from each other. Alleles from the cultivated zombi pea increased leaf width, while alleles from the wild zombi pea increased leaf length.

**Tuber traits** (NTB, TBW, TBL and TBWT): No QTL was detected for NTB and TBL. These two traits showed small trait variation and/or low heritability (27.7% for NTB) (Table 3). For TBW, five QTLs were identified on LGs 1, 2, 3, 4 and 8 in which the QTL on the LG8 had the largest effect (PVE=16.23%). In case of TBWT, three QTLs were detected on LGs 2, 4 and 8. Again, QTL on the LG8 showed the highest PVE (14.66%) and co-located with the QTL for TBW. At the QTLs detected on LGs 1, 3 and 8 alleles from the cultivated parent enhanced tuber size, while at the QTLs detected on LGs 2 and 4 alleles from the cultivated parent reduced tuber size.

**Plant type.** Stem length was considered.

**Stem length** (STL10): No QTL identified for this trait, although the trait showed high heritability (83.4%).

**Earliness.** Days to first flowering and days to first maturity were considered.

**Days to first flowering** (FLD). The cultivated and the wild parents were very different in FLD (102.6 and 72.4 days, respectively). The F_2_ population showed a high variation for the trait (Table 3). However, only one QTL was detected for this trait. The QTL was on LG1 and accounted for only 17.37% of the trait variation. The alleles from the cultivated parent prolong FLD.

**Days to first maturity** (PDDM). Similar to FLD, the cultivated and the wild parents were different in PDDM (34.0 and 27.8 days, respectively). Two QTLs on LG2, *Pddm2.1-* and *Pddm2.2+*, were identified for the trait. These QTLs showed similar PVE, 18.92% and 15.79%, respectively. The alleles from the cultivated parent at *Pddm2.1-* prolong PDDM, while that at *Pddm2.2+* hasten PDDM.

**Yield potential** or Number of seed per pod (SDDNPPD). The cultivated parent produced lower SDDNPPD than the wild parent. Five QTLs for SDDNPPD were detected LGs 5, 7, 8 and 9. Two QTLs on LG7 had high effects (PVE = 25.14% and 11.89%). At QTLs *Sddnppd5.1-* and *Sddnppd7.2-*, the allele from the cultivated parents increased SDDNPPD, while at the other three QTLs the alleles from the wild parent increased SDDNPPD.

**Pigmentation.** Seed coat color (SDC) and black mottle on seed coat (SDCBM)

**Seed coat color** (SDC): SDC was characterized as a qualitative trait. The cultivated parent had yellow seed coat but wild parent had brown seed coat color. F_2_ plants segregated for this trait at ratio of 81 (brown) to 24 (yellow), fitting a 3:1 ratio (χ^2^=0.26, *P*=0.61). This indicated that SDC is controlled by a single locus, named as *Sdc*. This locus was mapped as a morphological marker onto LG4 (Fig 2).

**Black mottle on seed coat** (SDCBM): SDCBM was also characterized as a qualitative trait. The cultivated Bali had no black mottle, while the wild parent had black mottle on seed coat. F_2_ progenies segregated for seed coat color at ratio of 76 (black mottle presence) to 29 (black mottle absence), fitting a 3:1 ratio (χ^2^=0.38, *P* = 0.54). This indicated that SDCBM is controlled by a single locus. The resulted indicated that the presence of SDCBM is controlled by a single dominant locus, named as *Sdcbm. Sdcbm* was mapped next to the locus *Sdc* at marker interval VR045 and C_13986_405837 (Fig 2).

### Distribution of domestication-related traits

QTLs with large effect (PVE > 20%) were found on four linkage groups, viz. LGs 4, 5 and 7 (Table 4).

**LG4**. Large-effect QTLs for pod length (*Pdl4.2+*) was on this linkage group. In addition, seed coat color (SDC) and black mottle (SDCBM) were mapped as morphological makers onto this linkage group.

**LG5.** Three large-effect QTLs were located to this linkage group; *Sdt5.1-* for seed thickness and *Lfpw5.1-* and *Lfmw5.1-* for leaf width. *Lfpw5.1-* and *Lfmw5.1-* with PVE = 34.03% and PVE = 52.16%, respectively were located closed together between markers VES0151 and CLM808. These two QTLs were likely the same locus. In addition, QTLs *Sdt5.1-*, *Pdl5.1+*, and *Sdnppd5.1-* were clustered together on this linkage group (Fig 4).

**LG7.** Most of large-effect QTLs for seed-related traits (*Sd100wt7.1-*, *Sdl7.1-* and *Sdw7.1-*) were located on this linkage group. These QTLs were clustered with QTLs for pod width (*Pdw7.1--*), number of seeds per pod (*Sdnppd7.1+*) and stem thickness (*Stt7.1-*). In addition, another QTL for number of seeds per pod (*Sdnppd7.2-*) was also located in this linkage group.

### Comparison of QTLs for domestication in *V. vexillata* with those of other *Vigna* species

The QTLs for domestication-related traits for zombi pea identified in this study were compared with those for yardlong bean [20], azuki bean [13], mungbean [16], and rice bean [17] based on common SSR markers (Fig 4). The comparison showed that most of the QTLs detected for zombi pea were also found in other *Vigna* species; they were mapped to the same linkage groups. For example, most of the QTLs for seed-related traits with high PVE in zombi pea were mapped onto LG7 in which several QTLs for seed-related traits were detected for yardlong bean, azuki bean, mungbean and rice bean. Three QTLs, *Pddm2.1-*, *Sdt8.1+* and *Sdnppd9.1-*, were mapped to similar regions with other *Vigna* species. Nonetheless, none or a few of QTLs for seed-related traits were detected on LG6 and LG10 of zombi pea which is the same case as in other *Vigna* species except yardlong bean.

## Discussion

### Fertility of progenies of cultivated zombi pea from Bali x wild zombi pea

A previous study on reproductive compatibility among cultivated zombi pea from Bali (tuber type zombi pea),cultivated zombi pea from Africa (seed type zombi pea), and wild zombi pea from Africa and Australia revealed no productive incompatibility between the African cultivated zombi pea (seed type zombi pea) and the wild form from both Africa and Australia, but revealed various pre- and post-zygotic compatibilities between the Bali cultivated zombi pea (seed type zombi pea) and the wild form from Africa and Australia [8]. However, in our present study, we successfully obtained an F_2_ population from a partially fertile F_1_ hybrid of a cross between the cultivated zombi pea from Bali using the former as female parent and wild zombi pea, this suggested that the Bali cultivated zombi pea constitutes a primary gene pool with the wild zombi pea. The difference results in our study and that in Damayanti et al. [8] is possibly due to environmental factors. Damayanti et al. [8] noted that even self-pollination of the cultivated zombi pea from Bali results in low pod setting. In our study, when we conducted the hybridization during November 2014 to February 2015, the Bali cultivated zombi pea showed very low pod setting (<3%). When it was grown again during November 2015 to February 2016 and during November 2016 to February 2017, it set more flowers and pods (~15% and ~70%, respectively; Somta and Dachapak, personal observation). In addition to environmental factors, genetic relatedness of the parents appeared to affect the successful hybridization of between different forms of zombi pea. In our study, the wild zombi pea (JP235863 from India) and the Bali cultivated zombi pea were genetically closely related [4], while in the study of Damayanti et al. [8] the wild zombi pea used were from Africa and Australia in which they were genetically highly differentiated from the Bali cultivated zombi pea [4].

### Transferability and polymorphism of SSR markers

Although as high as 1,876 SSR primers pairs from various *Vigna* species were screened for polymorphism between the cultivated and wild parents, only 36.6% of them were amplifiable and only 10.7% of them were polymorphic. The low polymorphism of SSR markers was also observed in the mapping parents of *V. vexillata* used by Marubodee et al. [33] in which only 6.2% of 1,336 SSR markers were polymorphic. In addition, Dachapak et al. [4] reported that only 21.2% of 1,024 SSR markers screened in 6 accessions of *V. vexillata* from Asia, Africa, Australia and America were polymorphic. However, the percentage of amplifiable SSR markers in Marubodee et al. [33] and Dachapak et al. [4] (65.4% and 58.1%, respectively) was higher than in our study (36.6%). It is worth noting that almost all of the markers used by Marubodee et al. [33] and Dachapak et al. [4] were used in our study. Nonetheless, these results indicate moderate transferability but low polymorphism of SSR markers from other *Vigna* species in *V. vexillata*. High transferability rate of SSRs (>80%) among several other *Vigna* species have been reported earlier [19,25,29,34,42]. The low transferability rate of SSRs from other *Vigna* species to *V. vexillata* is likely due to the fact that those *Vigna* species and *V. vexillata* are genetically highly different [43]. Due to the low transferability and polymorphism of SSR markers in *V. vexillata*, other types of markers (RAD-seq in this study) should be used for genome analysis of *V. vexillata*.

### Linkage map construction and chromosomal translocations in zombi pea

Previously, there was only two genetic linkage maps developed for zombi pea [33,44]. The first map contained 14 linkage groups with all were dominant markers (70 RAPD and 47 AFLP) except one co-dominant SSR marker. The map reported by Marubodee et al. [33] comprised 11 linkage groups from 84 SSR and 475 RAD-seq markers. The number of linkage map of zombi pea constructed in our study comprised 11 linkage groups of 145 SSR markers and 117 RAD-seq markers. The number of linkage groups of the maps developed by Marubodee et al. [33] and by this present study corresponded with the haploid chromosome number of *V. vexillata*. (2n = 2x = 22).

Previous comparative genome studies in the genus *Vigna* by means of comparison of linkage maps revealed high genome conservation among the species within this taxon [17,19,25,42,45] with exception for *V. vexillata* which several chromosome translocations occurred in this species; LG5, LG6 and LG9 were composed of the chromosome translocations from two linkage groups of mungbean or cowpea [33]. We compared our *V. vexillata* linkage map with other *Vigna* linkage maps (Fig 3). We found that middle to lower part LG5 of our *V. vexillata* linkage map was very likely a translocation from LG4 and that LG7 of our *V. vexillata* linkage map was translocated from LG4 and LG10. In addition, a part of the LG7 appeared to be duplication of LG2 or vice versa (Fig 3). Intraspecific macro-translocation has been demonstrated in azuki bean by means of comparative linkage mapping and fluorescence *in situ* hybridization [46,47]. The translocation was reciprocal-translocation where LG4 and LG6 were intermingled. This translocation was found in many accessions of wild azuki bean and it possibly affected fitness of those wild accessions to some environments and seed size [47]. In our study, comparative linkage analysis revealed macro-translocation between our zombi pea linkage map and the zombi pea linkage map developed by Marubodee et al. [33]; a part of LG10 of the latter translocated to LG7 of the former (Fig 3). This indicates intraspecific macro-translocation of zombi pea. However, it is not known whether this translocation provides any adaptive advantages for the zombi pea.

### Segregation distortion

Segregation distortion is a normal phenomenon in interspecific hybridization and in even intraspecific hybridization between two highly genetically different genotypes such as between wild and cultivated form of the same species. Segregation distortion of DNA markers has been reported for intraspecific hybridization of several *Vigna* species including *V. radiata* [16], *V. angularis* [41,46], *V. umbellata* [17], *V. mungo* [42] and *V. unguiculata* [19,20]. The percentage of marker distortion in these reports ranged from 3.9% in *V. angularis* to 48.7% in *V. unguiculata*. In this study, as high as 48.5% of the markers showed segregation distortion, the value is considered very high as compared to that other reports in *Vigna* species. The distorted markers were clustered on LGs 5, 6, 7, 8, 10 and 11 (Fig 2). The presence of a gene(s) controlling sterility and/or compatibility may cause segregation distortion of nearby loci and clustering of distorted markers [48]. The highly distorted markers (*P*< 0.001) were on LGs 5, 7 and 8. The distortions found on LG5 and LG7 possibly stem from the chromosomal translocations in these two linkage groups (Fig 2; see also above discussion on translocation). The cause of the distortion on LG8 is not known, however, it is possible that this linkage group represents the most genome divergence between the wild and cultivated parents used in this study. It is worth noting that major QTLs for tuber-related traits (*Tbw8.1-* and *Tbwt8.1-*) were located on the LG8 (Fig 4 and Table 4). Tuber root trait in the cultivated *V. vexillata* from Bali used in this study is the most strikingly different adaptive/domestication trait distinguishing the cultivated *V. vexillata* from Bali from other forms of *V. vexillata*. In yardlong bean (*V. unguiculata*), major QTLs for pod length and other pod-related traits (the most important domestication trait for this crop) and seed traits were found on LG7 where highly segregation distortions existed [19]. Similarly, in mungbean, major QTLs for seed dormancy, seed productivity and day length sensitivity were located on LG4 that showed high level of segregation distortion [16]. Most of markers mapped on LG11 of intra-and interspecific crosses of *Vigna* species always showed segregation distortion [17,19,45]. This is also the case in our study where 58.8% of the markers mapped to LG11 showed segregation distortion (Fig 2). Therefore, LG11 appeared to play an important role in genetic differentiation within and among *Vigna* species.

Although pod length (PDL) and number of seed per pod (SDNPPD) were highly correlated (*r* = 0.83) (S2 Table), only one of the QTLs for these two traits were co-located (Table 4 and Fig 4). Since three of the five QTLs, including the largest effect QTL, detected for SDNPPD were mapped to genome regions that showed highly segregation distortion, those QTLs may be associated with genetic compatibility and fertility, and thus seed production. In *Vigna* species, early generation (F_1_ and F_2_) hybrid progenies derived from hybridization of distantly-related genotypes always possesses pods with incomplete filled (empty seeds in some locules) [49,50] This may account for the no co-localization of QTLs for pod PDL and SDNPPD of *V. vexillata* in this study.

### QTLs for tuber weight in zombi pea

Most of edible tubers or storage roots of crop plants are from non-legume crops such as sweet potato, yam and cassava. In *Vigna* species, *V. vexillata, V. lobatifolia* and *V. marina* produced tuber roots [51]. Tubers produced by legume crops have much more nutrition than nonlegume crops especially nitrogen or protein due to nitrogen fixation ability of the legumes. The protein content in tuber roots of *V. vexillata* is about 15% which is five-fold higher than that in sweet potato. In our study, three QTLs were identified for tuber weight (TBWT) in *V. vexillata* with the QTL with largest PVE of about 15%. In potato, tuber weight was controlled by a few QTLs with PVE of 20% or less [52]. Thus, tuber weight *V. vexillata* and potato is controlled by small number of loci and is highly affected by environment. Tuber weight of potato has been shown to be negatively correlate with leaf area (leaf size) [53]. In our study, none of the QTLs for tuber weight co-located with QTLs for leaf size (Table 4 and Fig 4), indicating no genetic relationship between the two traits in *V. vexillata*. Among the three QTLs identified for tuber weight in *V. vexillata*, alleles from wild *V. vexillata* at two loci increased the tuber weight (Table 4). These alleles together the alleles for tuber weight from the Bali cultivated *V. vexillata* accounted for transgressive segregation found in the F_2_ population (S1 Fig). The alleles for tuber weight from the wild *V. vexillata* will be useful for improvement of tuber size in the Bali cultivated *V. vexillata*.

### Genomic region and distribution of QTLs for domestication traits

Many domestication-related traits were studied in *Vigna* crops including mungbean [16], yardlong bean [19,20], azuki bean [13] and rice bean [17]. In general, these studies found that (i) domestication-related traits were controlled by one or two major QTLs together with some minor QTLs in a narrow genome region, (ii) major QTLs for different traits were clustered and (iii) QTL clusters were not randomly distributed along linkage groups. For examples, major QTLs for seed size, pod size, and seed dormancy were clustered on LG4 of mungbean, and major QTLs for pod length, seed size, pod shattering, leaf size, and stem thickness were clustered on LG7 of yardlong bean. In this study, we did not found clusters of major QTLs for different traits in zombi pea (Fig 4 and Table 4). However, we found that major QTL for tuber weight on LG8 clustered with four minor QTLs for with pod size, seed size and tuber size (42.0-50.6cM on LG8; Fig 4). Clustering of QTLs may be resulted from pleiotropic effect or closely linked genes.

### Comparison of domestication QTLs between *V. vexillata* and other *Vigna* species

Only QTLs having phenotypic effect more than 20% were considered (Table 5) and were mainly compared with yardlong bean [20].

**Table 5. Comparison of major QTLs (PVE > 20%) detected for domestication in F_2_ population (AusTRCF66514 × JP235863) of *Vigna vexillata* with the major QTLs (PVE > 20%) detected for domestication in other *Vigna* species.**

The cultivated zombi pea used in this study and yardlong bean are believed to be have been domesticated in Asia, although their origin of species is believed to be in Africa [4,19]. Large-effect QTLs of yardlong bean were detected on four linkage groups (LGs 1, 3, 7 and 11), while large-effect QTLs of zombi pea were found on three linkage groups (LGs 4, 5 and 7). Although some major QTLs were located on LG7 of both species, distributions of QTLs on this linkage group were different. The LG7 of yardlong bean contained several large-effect QTLs for seed size, pod size, leaf size, and shattering, whereas the LG7 of zombi pea contain large-effect QTLs for only seed size and number of seeds per pod. The difference is no surprise due to the fact that zombi pea was primarily domesticated/selected for large tuber roots [54], whereas yardlong bean was principally domesticated/selected for long and soft immature pods [5].

**Seed weight**. The 100-seed weight of the cultivated zombi pea was 7.3 g, while the wild zombi pea was only 2.1 g. Two QTLs for this trait were located on LG2 and LG7 (the largest effect (PVE =28.19%) was on LG7). In yardlong bean, ten QTLs of 100-seed weight were detected on nine linkage groups with the highest QTL effect on LG7 (24.6%). Although the majors QTL for seed weight of zombi pea and yardlong bean were both located on the LG7, they appeared to be different because the QTL region in zombi pea was a translocation from LG10 as compared to yardlong bean and other *Vigna* species (Fig 3).

**Pod size**. Pod size of the cultivated zombi pea is not much larger longer than that of the wild zombi pea (94.3 vs. 85.6mm. for length and 5.7 vs. 3.6 mm. for width, respectively). Six QTLs for pod size in zombi pea was found on LGs 1, 2, 4, 5 and 7 with the largest effect QTL (PVE = 20.7%) on LG4. Seventeen QTLs for pod size in yardlong bean was found on every LGs except LG10 with the largest effect QTL on LG7 for both PDL and PDW (PVE = 31.0% and 31.2%, respectively). Some of QTLs related with pod size were located on same linkage in azuki bean. One of the QTL for pod size on LG4 was shared between zombi pea and yardlong bean (Fig 4) was found between markers CEDC055 and CEDG185.

**Leaf size**. Both primary and mature leaves of the cultivated zombi pea were larger than the wild zombi pea. Seven QTLs were detected for leaf size with highest effect on LG5 (*Lfpw5.1-*with PVE = 34.03% and *Lfmw5.1-*with PVE = 52.16%). In yardlong bean, thirteen QTLs for primary leaf were detected on LG1, LG2, LG3, LG6, LG7, LG8, LG9 and LG11, but only one of them (*Lfpl7.1+*) with highest PVE (20.9%) was found on LG7.

**Yield potential.** Number of seeds per pod (SDNPPD) of the cultivated zombi pea was lower than that of the wild zombi pea (7.6 and 12.7, respectively). Five QTLs were on LG5, LG7, LG8 and LG9 were detected with QTL showing highest PVE on LG7 (25.14%). In yardlong bean, two QTLs were detected on LGs 7 and 11. The QTL on LG11 showed highest PVE (70.1%).

### Comparison with azuki bean, mungbean and rice bean

Compared these eight QTLs (>20% PVE) of SD100WT, SDL, SDW, SDT, PDL, LFPW, LFMW and SDNPPD in zombi pea with QTLs domestication-related from azuki bean [13], mungbean [16] and rice bean [17].

**Seed size and weight.** Seven QTLs of seed size and weight were detected for zombi pea in which QTLs with high effect were on LG7 (*Sd100wt7.1-*, *Sdl7.1-* and*Sdw7.1-*). In azuki bean, rice bean and mungbean, large-effect QTLs for seed size and weight were co-located on the same region; LGs 1, 2, 3 and 9 for azuki bean, LG4 for rice bean, and LG8 for mungbean. This suggested that major genes controlling seed size with different biological mechanisms exist among domesticated *Vigna* species.

**Pod size.** Among the QTLs for pod length and pod width in zombi pea, the QTL *Pdl4.2+* on LG4 showed the highest PVE (20.65%). QTL with the highest effect (PVE = 23.1%) for pod length in rice bean was also detected on LG4, while the QTL with the highest effect for pod length in azuki bean and mungbean were on LG7.

**Leaf size.** Among seven QTLs for leaf size in zombi pea, two QTLs (*Lfpw5.1-* and *Lfmw5.1-*) showed PVE higher than 20%; both of which were on LG5. In rice bean, a major QTL for leaf size was located on LG4. No QTL was detected for leaf size in azuki bean and mungbean due to small difference in the mapping parents.

**Yield potential.** Five QTLs for SDNPPD in zombi pea were located on LGs 5, 7, 8 and 9 with the QTL on LG7 showed highest PVE (25.14%), while three QTLs for SDNPPD in rice bean were detected on LG4 and LG9, and two QTLs for SDNPPD in mungbean were identified on LG1 and LG9. The QTLs on LG7 of zombi pea, rice bean, and mungbean were identified in a similar location and may be the same locus. However, all the QTLs identified in rice bean and mungbean possessed PVE lower than 20%.

**Pigmentation.** Seed coat color and presence of black mottle on seed coat were treated as morphological markers which were mapped next to each other on the LG4. A QTL for pod length were located between the seed coat color locus and the black mottle locus. In azuki bean and mungbean, gene controlling the presence of black mottle on seed coat was also mapped on LG4. This indicated that the gene controlling this trait is highly conserved among species in the genus *Vigna*.

## Conclusions

An F_2_ of *V. vexillata* population derived from a partially fertile hybrid between tuber-root-type cultivated form and wild-type form were successfully developed. A genetic linkage map was constructed for this population utilizing 145 SSR, 117 RAD-seq and 2 morphological markers and to locate QTL for domestication syndrome traits. In total, 37 QTLs were detected for 18 domestication traits. Eight QTLs with large effect (>20%) were located on 4 out of 11 linkage groups. Domestication traits including seed size, pod size, leaf size, yield potential and seed pigmentation were controlled by one or two major QTLs and a few minor QTLs. QTLs for tubers were found on five different linkage group. Zombi pea had highest number of shared common QTLs with azuki bean, followed by yardlong bean, rice bean and mungbean. QTLs for seed size (SD100WT, SDL, SDW and SDT) were conserved and clustered together on different linkage groups of each species. These results provide a genetic map and information for marker-assisted selection of domestication-related QTLs in zombi pea and related species. The results also provide more understanding on genome evolution of *Vigna* species.

## Acknowledgement

This research was financially supported by Royal Golden Jubilee PhD programme (RGJ) co-funded by the Thailand Research Fund (TRF) and Kasetsart University.

## Supporting Information

**S1 Fig. Frequency distribution of 20 domestication related traits (except SDC and SDCBM traits) in F_2_ population of *Vigna vexillata* (AusTRCF66514 × JP235863).**

**S1 Table. Summary of percentage of amplification and polymorphic markers from four *Vigna* species in *Vigna vexillata* accessions AusTRCF66514 and JP235863.**

**S2 Table. Correlation between domestication traits in F_2_ population of *V. vexillata* (AusTRCF66514 × JP235863).**

## References

1. TomookaN, Kaga A, Isemura T, Vaughan DA, Srinives P, Somta P, et al. Vigna genetic resources. Proceeding of the 14^th^ NIAS international workshop on Genetic Resources, Genetic and Comparative Genomics of Legumes (Glycine and Vigna); 2010. pp. 11–21.

2. Tomooka N, Naito K, Kaga A, Saksai H, Isemura T, Ogiso-Tanaka E, et al. Evolution, domestication and neo-domestication of the genus Vigna. Plant Genet Resour 2014; 12(1): 168–171.

3. Ferguson H. The food crops of the Sudan and their relation to environment. Proceeding of Food and Society in the Sudan, Philosophical Society of the Sudan Khartoum, McCorquodale and Co. (Sudan) Ltd; 1954.

4. Dachapak S, Somta P, Poonchaivilaisak S, Yimram T and P Srinives. Genetic diversity and structure of the zombi pea (Vigna vexillata (L.) A. Rich) gene pool based on SSR marker analysis. Genetica 2017; 145: 189–200.

5. Karuniawan A, Iswandi A, Kale PR, Heinzemann J and WJ Grüneberg. Vigna vexillata (L.) A. Rich. cultivated as a root crop in Bali and Timor. Genet Resour Crop Evol 2006; 53(1): 213–217.

6. Bhattacharyya PK, Ghosh AK, Sanyal B and GD Ray. Grow Vigna vexillata for protein rich tuber cum pulse crop in the northern-eastern hill region. Seed Farms 1984. 10: 33–36.

7. Asati BS and DS Yadav. Diversity of horticultural crop in north eastern region. ENVIS Bull Himal Ecol 2004; 12: 1–11.

8. Damayanti F, Lawn RJ and LM Bielig. Genetic compatibility among domesticated and wild accessions of the tropical tuberous legume Vigna vexillata (L.) A. Rich. Crop Pasture Sci 2010; 61: 785–797.

9. Hammer K. Das Domestikationssyndrom. Genet Res Crop Evol 1984; 32: 11–34.

10. Gepts P. Domestication as a long-term selection experiment. Plant Breed Rev 2004; 24(2): 1–44.

11. Lush WM and LT Evans. The domestication and improvement of cowpea (Vigna unguiculata (L.) Walp.). Euphytica 1981; 30: 579–587.

12. Koinange EMK, Singh SP and P Gepts. Genetic control of the domestication syndrome in common bean. Crop Sci 1996; 36: 1037–1045.

13. Isemura T, Kaga A, Konishi S, Ando T, Tomooka N, Han OK, et al. Genome dissection of traits related to domestication in azuki bean (Vigna angularis) and their comparison with other warm season legumes. Ann Bot 2007; 100: 1053–1071.

14. Funatsuki H, Suzuki M, Hirose A, Inaba H, Yamada T, Hajika M, et al. Molecular basis of a shattering resistance boosting global dissemination of soybean. Proc Natl Acad Sci U.S.A. 2014; 111(50): 17797–17802.

15. Kaga A, Isemura T, Tomooka N and DA Vaughan. The genetics of domestication of the azuki bean (Vigna angularis). Genetics 2008; 178: 1013–1036.

16. Isemura T, Kaga A, Tabata S, Somta P, Srinives P, Shimizu T, et al. Construct of genetic linkage map and genetic analysis of domestication related traits in mungbean (Vigna radiata). PLoS One 2012; 7(8): e41304. doi:10.1371/journal.pone.0041304.

17. Isemura T, Kaga A, Tomooka N, Shimizu T and DA Vaughan. The genetics of domestication of rice bean, Vigna umbellata. Ann Bot 2010; 106: 927–944.

18. Andargie M, Pasquet RS, Muluvi GM and MP Timko. Quantitative trait loci analysis of flowering time related traits identified in recombinant inbred lines of cowpea (Vigna unguiculata). Genome 2013; 56(5): 289–294.

19. Kongjaimun A, Kaga A, Tomooka N, Somta P, Shimizu T, Shu Y, et al. An SSR-based linkage map of yardlong bean (Vigna unguiculata (L.) Walp. subsp. *unguiculata* Sesquipedalis group) and QTL analysis of pod length. Genome 2012; 55(2): 81–92.

20. Kongjaimun A, Kaga A, Tomooka N, Somta P, Vaughan DA and P Srinives. The genetics of domestication of yardlong bean, Vigna unguiculata (L.) Walp. ssp. unguiculata cv.-gr. sesquipedalis. Ann Bot 2012b; 109(6): 1185–2000.

21. Lo S, Munoz-Amatriain M, Boukar O, Herniter I, Cisse N, Guo YN, et al. Identification of genetic factors controlling domestication-related traits in cowpea (Vigna unguiculata L. Walp). bioRxiv 2017; 1–32. doi:10.1101/202044.

22. Garba M and RS Pasquet. The Vigna vexillata (L.) A. Rich. genepool. In: Sorensen M, Estrella JE, Hamann EOJ and SAR Ruiz, editors. Proceedings of the 2^nd^ International Symposium on Tuberous Legumes. 1996 Aug 5-8, Celaya, Guanajuato, Mexico; 1998. pp. 61–71.

23. Lodhi MA, Ye GN, Weeden NF and BI Reisch. A simple and efficient method for DNA extraction from grapevine cultivars and Vitis species. Plant Mol Biol Rep 1994; 12: 6–13.

24. Wang XW, Kaga A, Tomooka N, Vaughan DA. The development of SSR markers by a new method in plants and their application to gene flow studies in azuki bean [Vigna angularis (Wild.) Ohwi and Ohashi]. Theor Appl Genet 2004; 109: 352–360.

25. Chankaew S, Isemura T, Isobe S, Kaga A, Tomooka N, Somta P, et al. Detection of genome donor species of neglected tetraploid crop Vigna reflexo-pilosa (créole bean) and genetic structure of diploid species based on newly developed EST-SSR markers from azuki bean (Vigna angularis). PLoS One 2014; 9(8): e104990. doi:10.1371/journal.pone.0104990.

26. Li CD, Fatokun CA, Ubi B, Singh BB and GJ Scoles. Determining genetic similarities and relationships among cowpea breeding lines and cultivars by microsatellite markers. Crop Sci 2001; 41: 189–197.

27. Somta P, Musch W, Kongsamai B, Chanprame S, Nakasathien S, Toojinda T, et al. New microsatellite markers isolated from mungbean (Vigna radiata (L.) Wilczek). Mol Ecol Resour 2008; 8:1155–1157.

28. Seehalak W, Somta P, Sommanas W and P Srinives. Microsatellite markers for mungbean developed from sequence database. Mol Ecol Res 2009; 9: 862–864. doi:10.1111/j.1755-0998.2009.02655.x.

29. Tangphatsornruang S, Somta P, Uthaipaisanwong P, Chanprasert J, Sangsrakru D, Seehalak W, et al. Characterization of microsatellites and gene contents from genome shotgun sequences of mungbean (*Vigna radiata* (L.) Wilczek). BMC Plant Biol 2009; 9: 137. doi:10.1186/1471-2229-9-137

30. Yu K, Park SJ, Poysa V and P Gepts. Integration of simple sequence repeat (SSR) markers into a molecular linkage map of common bean (Phaseolus vulgaris L.). J Hered 2000; 91(6): 429–34.

31. Blair MW, Pedraza F, Buendia HF, Gaitan-Solis E, Beebe SE, Gepts P, et al. Development of genome-wide anchored microsatellite map for common bean (Phaseolus vulgaris L.). Theor Appl Genet 2003; 107(8): 1362–1374.

32. Hanai LR, de Campos T, Camargo LE, Benchimol LI, de Souza AP, Melotto M, et al. Development, characterization, and comparative analysis of polymorphism at common bean SSR loci isolated from genic and genomic sources. Genome 2007; 50(3): 266–27.

33. Marubodee R, Ogiso-Tanaka E, Isemura T, Chankaew S, Kaga A, Nalito K, et al. Construction of an SSR and RAD-marker based molecular linkage map of Vigna vexillata (L.) A. Rich. PLoS One 2015; 10(9): e0138942. doi:10.1371/journal.pone.0138942.

34. Somta P, Seehalak W and P Srinives. Development, characterization and cross-species amplification of mungbean (Vigna radiata) genic microsatellite markers. Conserv Genet 2009; 10:1939–1943.

35. Peterson BK, Weber JN, Kay EH, Fisher HS and HE Hoekstra. Double digest RADseq: an inexpensive method for de novo discovery and genotyping in model and non-model species. PLoS One 2012; 7(5): e37135. doi:10.1371/journal.pone.0037135

36. Catchen JM, Aemores A, Hohenlohe P, Cresko W and JH Postlethwait. Stack: building and genotyping loci de novo from short-read sequences. G3 (Bethesda) 2011; 1(3): 171–182.

37. Van Ooijien JW. 2006. Join Map 4.0, Software for the calculation of genetic linkage maps in experimental populations. Wageningen, Netherlands: Kyazma B. V.

38. Kosambi DD. The estimation of map distances from recombination values. Ann Eugenic 1944; 12: 172–175.

39. Li H, Ye G and J Wang. A modified algorithm for improvement of composite interval mapping. Genetics 2007; 175: 361–374.

40. Meng L, Li H, Zhang L and J Wang. QTL IciMapping: Integrated software for genetic linkage map construction and quantitative locus mapping in biparental populations. Crop J 2015; 3(3): 269–283.

41. Han OK, Kaga A, Isemura T, Tomooka N and DA Vaughan. A genetic linkage map for azuki bean (Vigna angularis). Theor App Genet 2005; 111: 1278–1287.

42. Chaitieng B, Kaga A, Tomooka N, Isemura T, Huroda Y and DA Vaughan. Development of black gram [Vigna mungo (L.) Hepper] linkage map and its comparison with an azuki bean [Vigna angularis (Wild.) Ohwi and Ohashi] linkage map. Theor Appl Genet 2006; 13(7): 1261–1269.

43. Takahashi Y, Somta P, Muto C, Iseki K, Naito K, Pandiyan M, et al. Novel genetic resources in the genus Vigna unveiled from gene bank accessions. PLoS One 2016; 11(1): e0147568. doi:10.1371/journal.pone.0147568

44. Ogundiwin EA, Thottappilly G, Aken’Ova ME, Ekpo EJA and CA Fatokun. Resistance to cowpea mottle carmovirus in Vigna vexillata. Plant Breeding 2002; 121: 517–520.

45. Somta P, Kaga A, Tomooka N, Kashiwaba K, Isemura T, Chaitieng B, et al. Development of an interspecific Vigna linkage map between Vigna umbellata (Thumb.) Ohwi & Ohashi and V. nakashimae (Ohwi) Ohwi and Ohashi and its use in analysis of bruchid resistance and comparative genomics. Plant Breeding 2006; 125(2): 77–84.

46. Kaga A, Isemura T, Tomooka N and DA Vaughan. The genetics of domestication of the azuki bean (Vigna angularis). Genetics 2008; 178: 1013–1036.

47. Wang L, Kikuchi S, Muto C, Naito K, Isemura I, Ishimoto M, et al. Reciprocal translocation identified in Vigna angularis dominates the wild population in East Japan. J Plant Res 2015; 128: 653–663.

48. Zamir D and Y Tadmor. Unequal segregation of nuclear gene in plants. Bot Gaz 1986; 147: 355–358.

49. Gosal SS and YPS Bajaj. Interspecific hybridization between Vigna mungo and Vigna radiata through embryo culture. Euphytica 1983; 32: 129–l37.

50. Gupta VP, Plaha P and PK Rathore. Partly fertile interspecific hybrid between a black gram x green gram derivative and an adzuki bean. Plant Breeding 2002; 121: 182–183.

51. National Research Council. Tropical legumes: resources for the future. The national Academies Press; 1979.

52. Rak K, Bethke PC and JP Palta. QTL mapping of potato chip color and tuber traits within an autotetraploid family. Mol Breed 2017; 37:15.

53. Lemaga B and K Caesar. Relationship between number of main stems and yield components of potato (Solanum tuberosum L. cv. Erntestolz) as influenced by different daylengths. Potato Res 1989; 33: 257–267.

54. Maréchal R, Mascherpa JM and F Stainier. Etude taxonomique d’un groupe d’especes des genres Phaseolus et Vigna (Papilonaceae) sur la base des donnees morphologiques et polliques, traitees pour l’analyse informatique. Boissiera 1978; 28: 1–273.

